# Functional dissection of Alzheimer’s disease brain gene expression signatures in humans and mouse models

**DOI:** 10.1101/506873

**Authors:** Ying-Wooi Wan, Rami Al-Ouran, Carl Grant Mangleburg, Tom V. Lee, Katherine Allison, Sarah Neuner, Catherine Kaczorowski, Vivek Phillip, Gareth Howell, Heidi Martini-Stoica, Hui Zheng, Jungwoo Wren Kim, Valina Dawson, Ted Dawson, Ping-Chieh Pao, Li-Huei Tsai, Jean-Vianney Haure-Mirande, Minghui Wang, Michelle E. Ehrlich, Hongkang Mei, Xiaoyan Zhong, Paramita Chakrabarty, Yona Levites, Todd E. Golde, Allan I. Levey, Accelerating Medicines Partnership-Alzheimer’s Disease Consortium, Benjamin Logsdon, Lara Mangravite, Zhandong Liu, Joshua M. Shulman

**Author notes:** Full consortium author list available at DOI: 10.7303/syn17114455. Co-first authors, contributed equally. Co-senior authors, contributed equally (Zhandong Liu, Joshua M. Shulman). Correspondence: Joshua M. Shulman, MD, PhD, Jan and Dan Duncan Neurological Research Institute, 1250 Moursund St., Suite N.1150, Houston, TX 77030, Zhandong Liu, PhD, Jan and Dan Duncan Neurological Research Institute, 1250 Moursund St., Suite N.1150, Houston, TX 77030, /.

## Abstract

Human brain transcriptomes can highlight biological pathways associated with Alzheimer’s disease (AD); however, challenges remain to link expression changes with causal triggers. We have examined 30 AD-associated, gene coexpression modules from human brains for overlap with 251 differentially-expressed gene sets from mouse brain RNA-sequencing experiments, including from models of AD and other neurodegenerative disorders. Human-mouse overlaps highlight responses to amyloid versus neurofibrillary tangle pathology and further reveal age- and sex-dependent expression signatures for AD progression. Human coexpression modules enriched for neuronal and/or microglial genes overlap broadly with signatures from mouse models of AD, Huntington’s disease, Amyotrophic Lateral Sclerosis, and also aging. Several human AD coexpression modules, including those implicated in the unfolded protein response and oxidative phosphorylation, were not activated in AD models, but instead were detected following other, unexpected mouse genetic manipulations. Our results comprise a powerful, cross-species resource and pinpoint experimental models for diverse features of AD pathophysiology from human brain transcriptomes.

## INTRODUCTION

Alzheimer’s Disease (AD) is a progressive and incurable neurodegenerative disorder, with rapidly increasing prevalence due to population aging (Scheltens et al., 2016). At autopsy, AD is characterized by extracellular neuritic amyloid plaques and intraneuronal neurofibrillary tangles, comprised of misfolded and aggregated Amyloid-Beta (Aβ) peptide and the Microtubule Associated Protein Tau (MAPT/Tau), respectively. Based on RNA sequencing (RNA-Seq) from 2,114 human postmortem brain samples, the Accelerating Medicines Partnership-Alzheimer’s disease (AMP-AD) consortium has identified 30 coexpression modules significantly associated with AD clinical-pathologic diagnosis (Logsdon et al., submitted). These data show promise to highlight completely unexpected molecular insights into AD pathogenesis; however, interpretation is hindered by several obstacles. One major challenge arises from the recognition that the AD pathologic cascade initiates decades prior to onset of clinical manifestations (De Strooper et al., 2016; Jack et al., 2013), whereas human brain expression profiles can only be generated cross-sectionally at the time of death. Indeed, it is essential to reconstruct the longitudinal, aging-dependent time-course of molecular derangements in order to pinpoint the earliest possible opportunities for intervention and to develop effective biomarkers. Second, most brains from older persons with AD show mixed pathologies at autopsy, including many other age-related lesions associated with cognitive impairment (Kapasi et al., 2017). Therefore, functional dissection of human coexpression modules requires differentiating specific causal triggers, including (i) AD lesions (Aß plaques and/or Tau tangles), (ii) other neuropathology, or (iii) brain aging more generally. Lastly, even following the most comprehensive analysis, it remains challenging to determine which gene expression changes associated with disease are truly primary and therefore causal, rather than simply a consequence of AD.

By contrast with studies of human postmortem tissue, mouse models allow for controlled experimental manipulations to isolate the effects of specific molecular lesions, investigate for dynamic, age-dependent changes, and definitively establish causation. A large number of AD mouse models have been extensively characterized, and these systems have already contributed enormously to our understanding of disease pathogenesis (Götz and Ittner, 2008; Jankowsky and Zheng, 2017; LaFerla and Green, 2012). The most widely used transgenic models express mutant forms of the *amyloid precursor protein* (APP) gene, with or without *presenilin-1/2* (*PSEN1/2*), which are associated with autosomal dominant, early-onset AD; or alternatively, *MAPT* mutations, which cause familial frontotemporal dementia (FTD). These models recapitulate features of AD neuropathology, including plaques or tangles, along with variable degree of neuronal dysfunction/loss, and progressive neurobehavioral impairments (Ballatore et al., 2007; Esquerda-Canals et al., 2017). More recently, gene expression profiling, including RNA-seq, has been applied to elucidate brain transcriptome signatures. Among the results, these investigations have highlighted the importance of immune/inflammatory and neuronal/synaptic changes (Boisvert et al., 2018; Cummings et al., 2015; Gjoneska et al., 2015; Matarin et al., 2015; Rothman et al, 2018; Stephenson et al., 2018; Swartzlander et al., 2018). While some reports have identified selected overlaps between expression changes in AD mouse models and human brains (Bennett et al., 2018a; Castillo et al., 2017; Mostafavi et al., 2018; Neuner et al., 2019; Rojo et al., 2017), other studies have questioned the overall degree of conservation (Burns et al., 2015; Galatro et al., 2017; Hargis and Blalock, 2017).

The goal of this study is to systematically define correspondences between gene expression changes associated with AD in human brains and those caused by controlled experimental manipulations in mouse models. In addition to the brain RNA-seq datasets available from mouse models of AD, our cross-species analysis takes advantage of hundreds of other experimental comparisons relevant to diverse neurologic disorders, brain health, and aging. Our results highlight strengths and limitations among currently available AD models, and enhance understanding of the causes and consequences for gene expression signatures in human brains.

## RESULTS

To enable cross-species comparisons of brain transcriptome data, we reprocessed brain RNA-Seq data from 96 distinct mouse studies relevant to AD, other neurodegenerative disorders, aging and related mechanisms (Figure 1 and Table S2, Methods), and curated 376 unique experimental comparisons (genetic manipulation vs. control condition). The resulting 251 sets of significant DEGs (gene membership >10, fold-change > 1.2, FDR < 1%) define brain “gene expression signatures” (mean = 1385 genes, range = 10 – 12,393) characteristic of the respective mouse model comparisons (Table S3 and Supplemental Files). These expression profiles encompass 25,181 unique genes out of 52,873 total in the mouse transcriptome. As visualized using the t-SNE algorithm (Figure 1C), the curated expression signatures capture a combination of disease-specific and overlapping features characteristic of the heterogeneous mouse models included in this study. Even among mouse models for the same disease category, the overall extent of gene sharing among the corresponding sets of DEGs was modest (Figure S1); therefore, the included comparisons sample a wide spectrum of brain transcriptional responses. We next evaluated the overlap between each mouse expression signature and 30 human AD-associated brain coexpression modules, based on analyses of human postmortem brain RNA-seq from AMP-AD (Table S1) (Logsdon et al, submitted). Overall, we detect 1,569 significant overlaps (p_adj_ < 0.01), with the majority (68%) of experimental models showing evidence for gene expression changes corresponding to multiple human modules (mean = 6 overlapping modules per mouse model DEG set) (Figure 2). As expected, since human coexpression modules derived from different brain regions demonstrate extensive gene sharing, many mouse DEG sets show consistent overlaps within “module clusters” consisting of overlapping gene sets similarly enriched for neuronal, microglial, astrocytic, oligodendroglial, and/or endothelial expression signatures (Logsdon et al., submitted). We next assessed the direction of gene expression changes in mouse and human brains. The majority (77%) of overlapping gene expression signatures are concordant across species (Figure 2). For example, module clusters enriched for microglial genes are up-regulated in brain transcriptomes from human AD and mouse models, whereas those enriched for neuronal genes are predominantly down-regulated, but with notable exceptions (Figure 2 & Figure 3C). Lastly, 28 out of 30 modules significantly overlap (p_adj_ < 0.01) with at least 1 mouse model expression signature. In sum, these results support broad conservation of gene regulatory systems in the mammalian brain, and highlight that many AD-associated brain expression patterns are recapitulated in available mouse experimental models.

**Figure 1:**
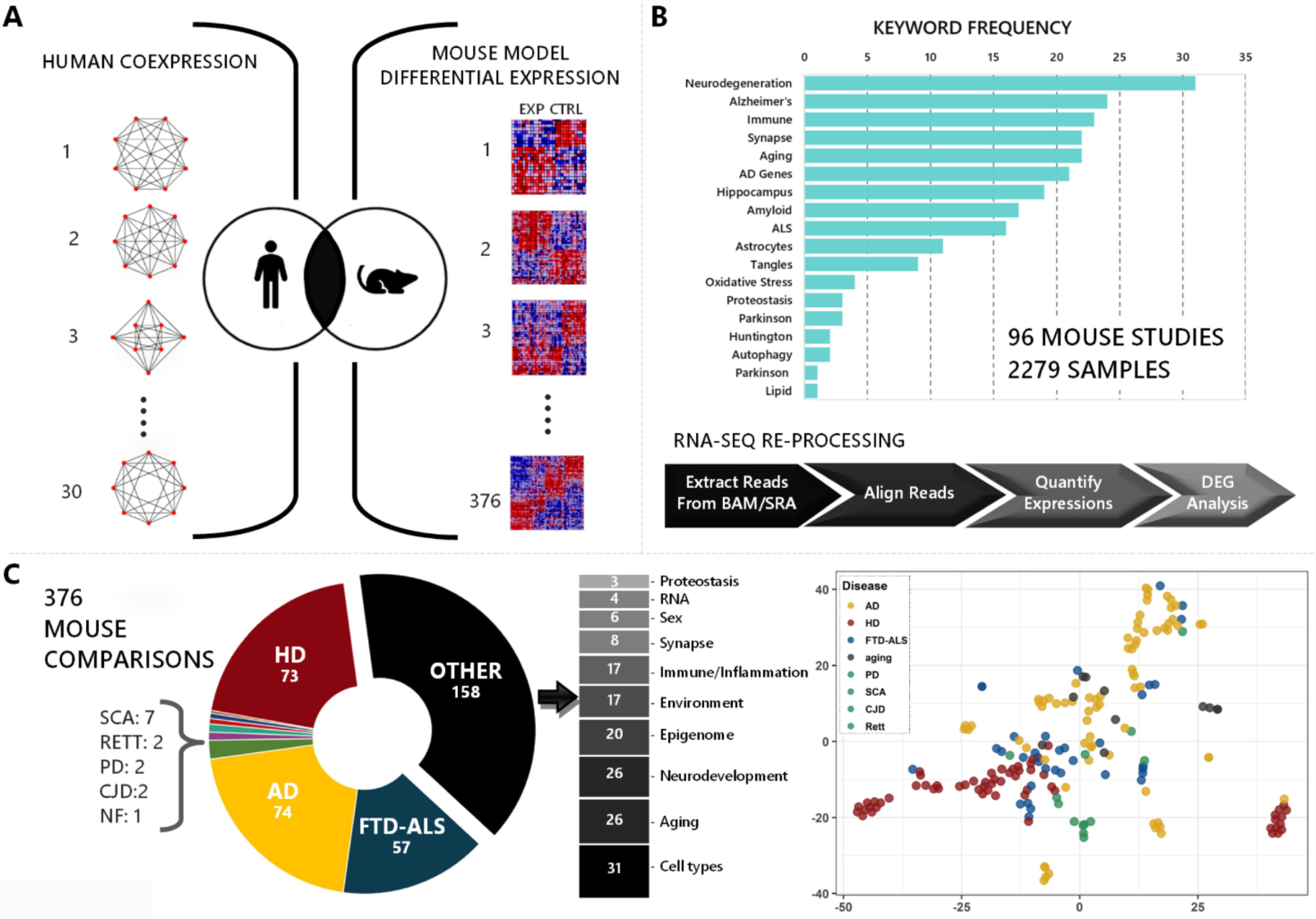
Study design and data. (A) We systematically examined overlaps between 30 Alzheimer’s disease (AD)-associated human co-expression modules (rows) and differentially-expressed gene (DEG) sets from 376 curated experimental comparisons in mouse models with RNA-seq profiles. (B) 96 mouse studies were selected based on relevance to AD, other neurodegenerative disorders, and implicated mechanisms. Distribution of keywords is shown among all included studies. All RNA-seq data was re-processed using a standard pipeline, and experimental versus control comparisons were manually reviewed and curated prior to differential expression analysis. (C) The 376 sets of DEGs considered in our analyses highlight expression signatures for mouse models of AD and other neurodegenerative disorders, including Huntington’s disease (HD), Frontotemporal Dementia-Amyotrophic Lateral Sclerosis (FTD-ALS), spinocerebellar ataxia 1 (SCA), Rett syndrome (RETT), Parkinson’s disease (PD), Creutzfeld Jacob disease (CJD), and neurofibromatosis (NF). We also considered DEGs from additional mouse models relevant to neurodegenerative mechanisms (Other). A T-distributed Stochastic Neighbor Embedding (t-SNE) plot was generated from all mouse model differential expression signatures included in this study, highlighting that these DEG sets capture a combination of disease-specific and overlapping features among heterogeneous neurodegenerative models. See also Figure S1 and S3.

**Figure 2:**
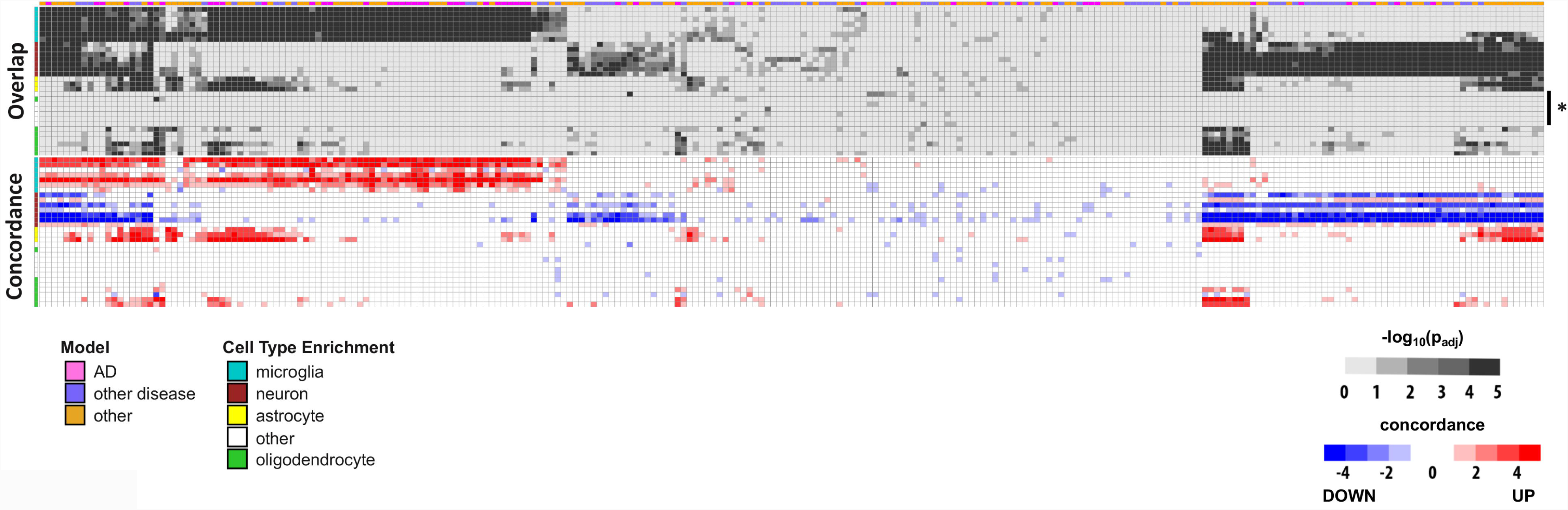
Overview of human-mouse overlaps and concordance. (A) Heat maps show overlap (top) and concordance (bottom) among 30 human coexpression modules (rows) and 251 sets of differentially expressed genes (DEGs, columns) from mouse model experimental comparisons. The average sample size for each comparison is 8.4 (range=4-28 total samples). Mouse-human overlap significance, calculated using the hypergeometric test, is represented in grayscale [-log_10_(p_adj_)]. Direction (red/blue) and extent of concordance (intensity) for gene expression changes in human brains and mouse models is also indicated (bottom). The color bar at top annotates all DEGs based on whether they derive from Alzheimer’s disease (AD) models (pink), other neurodegenerative disease models (purple), or other experimental manipulations potentially relevant to AD mechanisms (orange). The color bar at left denotes cluster membership for each human coexpression module, based on enrichment for cell-type specific gene signatures, including microglia (turquoise), neuron (brown), astrocyte (yellow), or oligodendrocyte (green). Module clusters associated with cell-type expression signatures (microglia, neuron, astrocytes, and oligodendrocytes) show broad overlaps with mouse model expression signatures. By contrast, the remaining modules poorly enriched for known cell type signatures (asterisk, right) show sparse overlap with mouse DEGs. Comprehensive results and details for all models and resulting overlaps can be found in Table S3 and S4.

**Figure 3:**
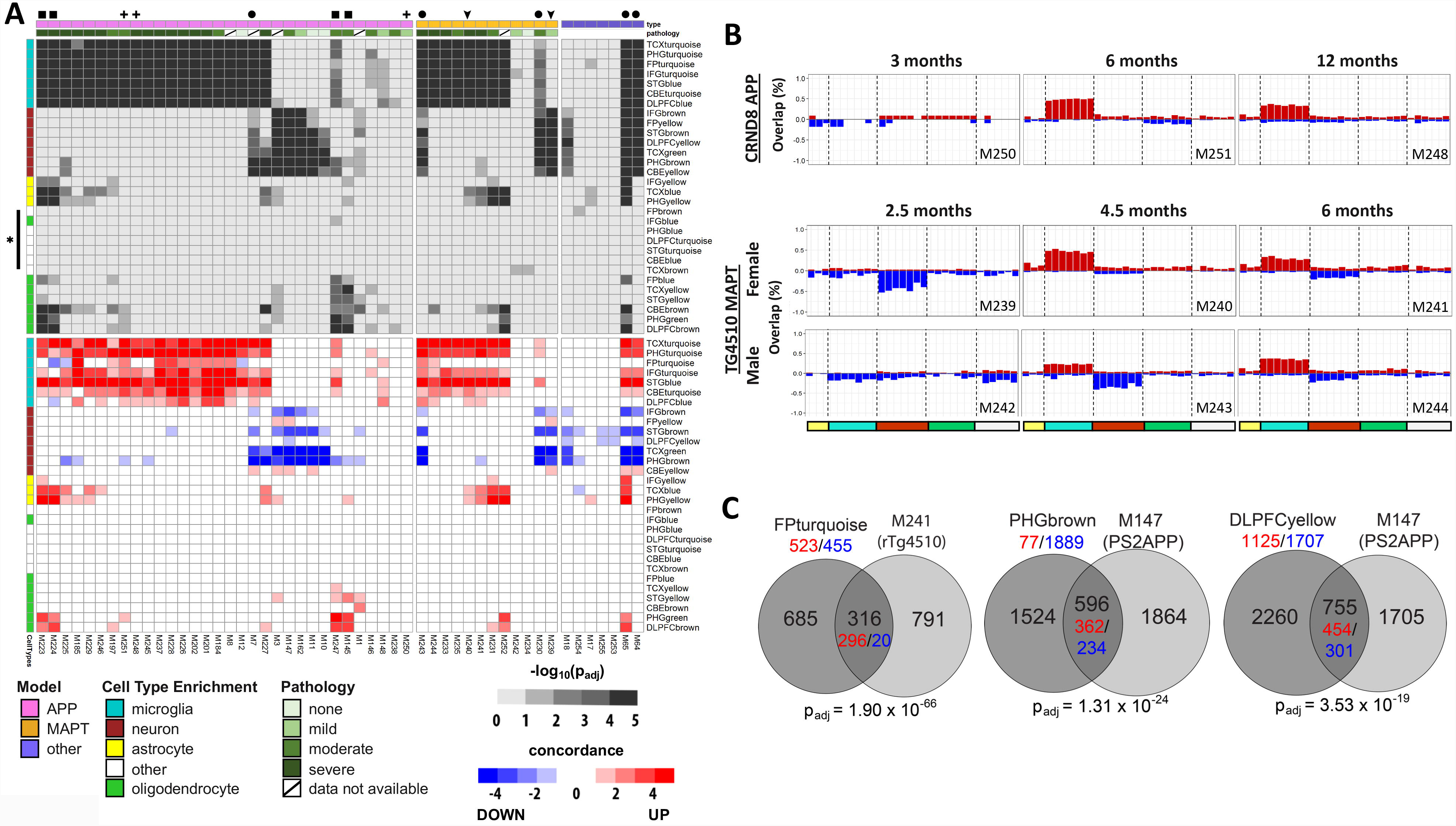
Human coexpression module overlaps with AD mouse models. (**A**) Heat maps show overlap (top) and concordance (bottom) among 30 human coexpression modules and sets of differentially expressed genes (DEGs) derived from Alzheimer’s disease (AD) mouse models. Mouse-human overlap significance, calculated using the hypergeometric test, is represented in grayscale [-log_10_(p_adj_)]. Direction (red/blue) and extent of concordance (intensity) for gene expression changes in human brains and mouse models is also indicated. The color bar at top annotates AD model comparisons based on whether they derive from APP (pink), MAPT (orange), or other (purple) AD models. The estimated pathologic burden (plaques/tangles and neuronal loss) in APP and MAPT models is also annotated in green, based on Alzforum (http://www.alzforum.org/research-models/). The color bars at left denotes cluster membership for each human coexpression module, based on enrichment for cell-type specific gene signatures, including microglia (turquoise), neuron (brown), astrocyte (yellow), or oligodendrocyte (green). The remaining modules poorly enriched for known cell type signatures (white, asterisk) show sparse overlap with DEGs from AD mouse models. Comprehensive results and details for all models and resulting overlaps can be found in the Supplemental Tables, cross-referenced by the unique ID code (bottom) for each set of model DEGs. Selected human module-mouse DEG overlaps referred to in the text are denoted as follows: ΔΔ, APP models with oligodendrocyte-enriched module overlaps; **+**, CRND8-APP models showing sustained, activation of microglial modules from 6 months onward; ▾, transient activation of neuronal modules in TG4510-MAPT model preceding microglial module overlap; •, co-activation of neuronal and microglial modules. Comprehensive results and details for all models, including sample sizes for all comparisons, and resulting overlaps can be found in Table S3 and S4. (**B**) Mouse model overlaps highlight age- and sex-dependent changes in the AD transcriptome. Direction of gene expression changes denoted by red (up) and blue (down). Magnitude of gene expression changes shown as percentage of the mouse DEG set overlapping each module. Cell type module clusters are denoted by colors on the bottom of the panel, as in A. (**C**) Representative overlaps of human modules with mouse DEGs. Gene counts are noted in black, including for overlapping and non-overlapping regions. To assess concordance between human brains and mouse models, gene counts are shown, noting increased or decreased expression (red or blue, respectively), including for the whole human coexpression module and the overlapping gene set from mouse models. See also Figure S2.

We next systematically examined whether available mouse models of AD show gene expression changes similar to those detected in human AD brains (Figure 3A). Our reprocessed dataset includes 53 expression signatures from 12 distinct AD transgenic models, including numerous *APP* and *MAPT* transgenic strains. Human microglial- and neuronal-enriched coexpression modules strongly overlap with expression signatures from mouse AD models (Figure 3A-C). For example, the microglial-enriched FPturquoise module overlaps significantly (p_adj_ < 1×10^−5^) with the majority of expression signatures from both *APP* (66%) and *MAPT* (67%) mouse models. While overlaps with the neuronal-enriched modules, such as PHGbrown, were more restricted, this pattern was still observed in a substantial proportion of models (22% and 25% of *APP* and *MAPT* models, respectively). In most cases, the attenuation of gene expression from neuronal modules and activation of microglial-enriched signatures appears to be mutually exclusive. Interestingly, coincident changes in both neuronal and microglial modules (Figure 3A, circles) was characteristic of selected *MAPT* transgenic strains, and was also seen in the *CDK5-P25* mouse which similarly develops extensive Tau pathology and brain atrophy (Cruz et al., 2003). This coincident microglial/neuronal overlap pattern is only rarely seen among expression signatures from the *APP* models (1 out of 34 DEG sets). We also considered overlaps in relation to the level of pathologic burden previously reported in each *APP* or *MAPT* model (Figure 3A). Overall, mouse expression signatures exhibited strong overlaps with microglial-enriched human modules at later, more severe stages of brain pathology. Reciprocally, neuronal module overlaps characterized models with earlier and comparatively mild pathologic burden. RNA-seq profiles generated from multiple aged timepoints permit examination of dynamic changes in human overlap patterns for selected AD models (Figure 3B). For example, in the TgCRND8 *APP* transgenic mouse, microglial module activation is seen by 6-months, and this signature is sustained in evaluations at 12-months (M248, M250, and M251; Figure 3A, cross-hatches and Figure 3B, top). By contrast, in the rTg4510 *MAPT* mouse model, transient gene expression changes overlapping neuronal-enriched human modules either preceded or accompanied the appearance of microglial expression signatures (M239-M244; Figure 3A, arrowheads, and Figure 3B, bottom). In addition, when compared to males, female rTg4510 mice revealed accelerated progression in brain transcriptional changes (Figure 3B, bottom), consistent with a sex by age interaction. Compared with the microglial- and neuronal-enriched module clusters, overlaps between AD mouse model expression signatures and astrocytic, oligodendroglial, and/or endothelial-enriched modules were more sparsely detected (Figure 3A). For example, TCXyellow and STGyellow—two modules similarly enriched for oligodendroglial markers and implicated in sphingolipid metabolism—showed selective overlap with aged *APP* transgenic models (e.g. TgCRND8 (M247), 5xFAD (M223), APPPS1 (M3) and PS2APP (M145); Figure 3A, squares, and Figure S2). In sum, our cross-species analyses highlight significant overlaps between transcriptional responses in AD mouse models and human postmortem brains, and pinpoint those expression signatures denoting age- and sex-dependent disease progression as well as Aß-versus Tau-pathology.

Numerous mechanistic parallels have been recognized between AD and other neurodegenerative disorders, including similar protein aggregate pathologies, proteostatic and oxidative stress, neuroinflammation, and the critical role of aging in disease risk and/or progression (Block and Hong, 2005; Bucciantini et al., 2002; Guo and Lee, 2014; Haass and Selkoe, 2007; Nixon, 2005; Ross and Poirer, 2004). We therefore examined potential overlap between human AD coexpression modules and differential expression signatures from other mouse experimental disease models, including Huntington’s disease (HD), frontotemporal dementia-amyotrophic lateral sclerosis (FTD-ALS), Parkinson’s disease, spinocerebellar ataxia (SCA), Creutzfeldt-Jakob disease (CJD) and Rett syndrome, along with available data from aged, wildtype mice mouse strains (Figure 4 and Figure S4-6). Similar to APP/MAPT transgenic mice, most disease models strongly activate expression signatures overlapping with neuronal- and microglial-enriched human modules. For example, the neuronal-enriched PHGbrown module overlaps with mouse brain expression signatures from models of HD (M94, p_adj_=2.5−10^−10^), FTD-ALS (*FUS*) (M173, p_adj_=4.8×10^−6^), SCA (M155, p_adj_=1.5×10^−18^), and prion disease (M200, p_adj_=6.0×10^−32^). Similarly, the microglial-enriched FPturquoise module significantly overlaps with models of FTD-ALS (*TDP43*) (M24, p_adj_=2.0×10^−42^), prion disease (M199, p_adj_=3.7×10^−18^), and Rett Syndrome (M198, p_adj_=3.1×10^−3^), and more selectively in HD (M81, p_adj_=1.1×10^−4^) and SCA (M151, p_adj_=5.3×10^−5^) models. Therefore, these signatures likely represent common brain transcriptome responses induced by diverse neurodegenerative triggers (e.g. APP, MAPT, Huntingtin, SCA1, and others). Consistent with this, we found that genes implicated in immune biology and inflammation comprise those recurring most frequently among the 251 mouse expression signatures included in this study (Figure S3). In addition, expression signatures from aged, wildtype mice also show significant overlaps with either neuronal- (M194) or microglial-enriched (M56, M219) human coexpression modules, or both (M220, M221) (Figure 4). For example, compared with 3-month controls, hippocampal tissue from 24-month-old mice (M56) revealed expression signatures overlapping with FPturquoise (p_adj_=2.9×10^−17^) and other modules implicated in inflammation and immunity. As discussed below, this result suggests that these patterns may not be specific for disease states but rather may accompany brain aging more generally. We therefore examined for any human coexpression modules with overlap profiles showing relative specificity for AD models. While none show absolute specificity, we found that modules enriched for oligodendroglial markers, including FPblue and TCXyellow are strongly activated in selected AD models (e.g. M145, M247, M223, M65 in Figure 3) but show comparatively sparse or weaker overlap with expression signatures from the majority of other disease models. Thus, these modules may highlight transcriptional programs that are preferentially activated by AD pathophysiology, such as Aß neurotoxicity.

**Figure 4:**
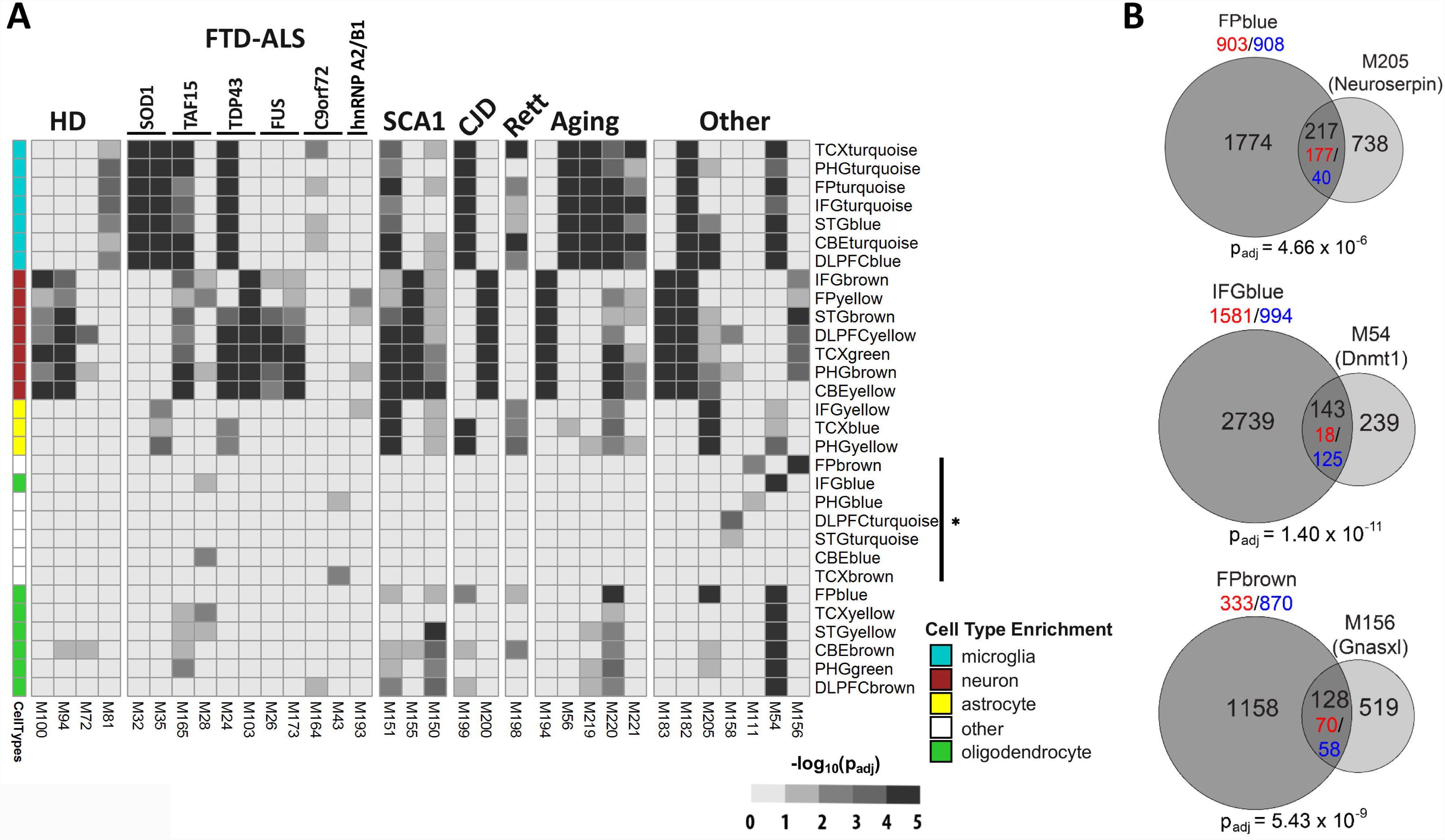
Overlaps with other mouse models. (**A**) Heat maps show overlap among human coexpression modules and sets of differentially expressed genes (DEGs) derived from mouse models, including pure aging, additional neurodegenerative disorders, and other experimental manipulations relevant to Alzheimer’s disease mechanisms. Mouse-human overlap significance, calculated using the hypergeometric test, is represented in grayscale [-log_10_(p_adj_)]. HD, Huntington’s disease; FTD-ALS, frontotemporal dementia-amyotrophic lateral sclerosis; SCA1, spinocerebellar ataxia 1; CJD, Creutzfeld-Jacob disease. Overlaps between HD model expression signatures and neuronal gene-enriched human coexpression modules recapitulate polyglutamine length- (M100, Q92 vs. M94, Q175) and brain region dependence (M100/M94, striatum vs. M72/M81, cortex). Other genetic manipulations generate expression signatures similar to AD mouse models, including (i) selective activation of microglial-enriched modules (M183, *PTCH1* knockout), (ii) coincident overlaps with microglial- and neuronal-enriched modules (M182, *nmf205*), or (iii) overlap with astrocytic- and oligodendroglial-enriched modules (M205, Neuroserpin). Modules poorly enriched for cell type signatures (denoted by asterisk, right) show sparse overlap with DEGs from AD mouse models, but selected overlaps are identified with FTD-ALS models (M28, M43) and other, unexpected genetic manipulations (M158, M111, M54, and M156). Comprehensive results and details for all models, including sample sizes for all comparisons, and resulting overlaps can be found in Table S3 and S4. For complete heat maps representing overlaps with HD, FTD-ALS, SCA1, and aging models, see Figure S4-6. (**B**) Representative overlaps of human modules with mouse DEGs, as in Figure 3C.

Our analyses also considered many additional mouse genetic manipulations based on potential relevance to neurodegenerative mechanisms (Figure 1B-C). Indeed, several other mice showed strong overlaps with human AD coexpression modules similar to canonical AD transgenic models (Figure 4A-B). For example, the C57BL6/J (B6J)-*nmf205* mouse (M182) has an expression signature which overlaps significantly with both the neuronal- and microglial-enriched human AD coexpression modules (PHGbrown, p_adj_=9.6×10^−39^, and FPturquoise, p_adj_=6.4×10^−6^, respectively), similar to that seen in selected *MAPT* transgenics (M230, M243) and the *CDK-P25* mouse (M64, M65). This mouse model consists of mutations in both the translation GTPase *GTPBP2* and the neuronal tRNA^Arg^_UCU_ gene causing neurodegeneration in association with ribosome stalling (Ishimura et al., 2014; Ishimura et al., 2016). In another example, expression of a mutant form of *neuroserpin* in mouse neural progenitor cells (Guadango et al., 2017) activates a transcriptional signature (M205) that significantly overlaps with the astrocyte-enriched modules, including IFGyellow (p_adj_=8.1×10^−17^), as well as FPblue (Figure 4B, p_adj_=4.7×10^−6^), an oligodendroglial-enriched module showing some selectivity for AD transgenic models (above). Autosomal dominant mutations in neuroserpin cause familial encephalopathy with neuroserpin inclusion bodies, a rare, young-onset neurodegenerative dementia caused by protein aggregation within the endoplasmic reticulum and associated ER stress (Roussel et al., 2013).

Of the 30 human AD-associated coexpression modules, a minority show virtually no overlap with AD mouse model brain expression signatures (asterisk, Figures 2,3A,4A). Such modules are strong candidates for features of AD pathobiology that are poorly recapitulated by existing AD mouse models. Interestingly, our cross-species analyses pinpoint a number of other mouse model expression signatures that overlap these modules (Figure 4A-B). For example, conditional knockout of the DNA methyltransferase, *DNMT1* (M54), in the mouse brain (Narayanan et al., 2014) activates a set of DEGs that significantly overlap (Figure 4B, p_adj_=1.4×10^−11^) with a human module, IFGblue, implicated in the unfolded protein response and DNA repair pathways. In another example, brain RNA-seq from a *Gnasxl*-deficient mouse (Holmes et al., 2016) (M156) overlapped with the FPbrown coexpression module (Figure 4B, p_adj_=5.4×10^−9^) that is enriched for genes involved in oxidative phosphorylation and mitochondrial translation. Interestingly, *Gnas*, which encodes a G-protein alpha stimulatory subunit, is a complex, imprinted genomic locus implicated in hypothalamic control of energy balance. Loss of the *Gnasxl* isoform causes a hypermetabolic mouse phenotype, resulting in growth retardation, hypoglycemia, and reduced adiposity (Nunn et al., 2013).

## DISCUSSION

Mouse genetic models have contributed enormously to our understanding of AD pathophysiology (Ballatore et al., 2007; Esquerda-Canals et al., 2017); however, the utility of these mice as robust preclinical models for AD has been challenged (Drummond and Wisniewski, 2017; Onos et al., 2016; Sasaguri et al., 2017). First, most AD mouse models are based on rare forms of familial autosomal dominant AD, which are caused by single, highly penetrant gene mutations. By contrast, late-onset AD arises from dozens of other risk variants, including many with modest effect sizes (Karch et al., 2014; Lambert et al., 2013; Sims et al., 2016), perhaps in combination with non-genetic risk factors. Second, unlike mouse models, most brain autopsies from individuals with AD show evidence of heterogeneous, mixed pathologies which likely modify disease onset, manifestations, and progression. Third, it has been suggested that widely used mouse neurobehavioral assays may be poor predictors of clinically relevant outcomes in human AD. In this study, we use the brain transcriptome to examine the correspondence between 30 human AD brain gene expression networks and 251 mouse experimental comparisons. These analyses provide a systems-level view of the molecular overlap between AD in humans and mouse models, as represented in the brain transcriptome.

Our results show that many AD transgenic mice, including both *APP* and *MAPT* models, show differential expression signatures that significantly overlap with AD-associated coexpression modules from human brains. The most robust overlaps were detected among module clusters strongly enriched for neuronal and microglial genes. However, these modules account for only 14 out of 30 total coexpression networks (47%), and as discussed further below they do not appear to be specific for AD. By contrast, a second group of human modules, especially those enriched for oligodendroglial genes, show comparatively greater specificity for AD, albeit for a rather restricted subset of models. Lastly, a substantial minority of AD-associated human brain coexpression modules had virtually no detectable overlap with available AD mouse models. The overlaps we define highlight those molecular features of AD biology recapitulated in existing AD mouse models, whereas non-overlapping modules may identify aspects of AD pathophysiology poorly captured. We conclude that most AD mouse models show overall poor correspondence to human disease, based on brain transcriptomes, with exception of neuronal/microglial-enriched modules. This is an important caveat for interpretation of studies using these animal models, and may explain in part their poor predictive power as AD preclinical models. Our findings are consistent with a complementary study of mouse data from gene expression arrays (Hargis and Blalock, 2017). Since the overall correlation of the brain transcriptome and proteome is modest (Seyfried et al., 2017), it will be important in future work to consider whether human-mouse gene expression overlaps are improved at the protein level.

Our analyses also highlight the value of experimental models for interpretation of human brain transcriptome profiles. Analyses considering either cross-sectional or longitudinal datasets similarly suggest that transcriptional changes overlapping human brain neuronal-enriched modules may represent an earlier, transient stage of AD, whereas microglial signatures are a subsequent and more sustained AD endophenotype. These results are consistent with other studies of AD mouse model transcriptomes reporting changes in neuronal- and microglial-enriched gene pathways (Cummings et al., 2015; Gjoneska et al., 2015; Matarin et al., 2015). Additionally, human brain astrocytic and oligodendroglial modules overlap with differential expression signatures from CDK5-P25 and 14 month-old 5xFAD models, which are both notable for their advanced pathologic changes, potentially consistent with a comparatively end-stage AD transcriptional endophenotype. Transition points between human-mouse overlaps can be linked to the manifestation of disease-relevant mouse phenotypes. For example, CRND8 *APP* mice manifest reduced synaptic markers and hippocampal neuronal loss at 6 months, when overlaps are first detected with microglial modules (Adalbert et al., 2009; Brautigam et al., 2012). Similarly, in Tg4510 *MAPT* transgenic animals evaluated at 4 and 6 months, respectively, memory task impairment and subsequent neurodegenerative pathologic changes correspond to sequential activation of neuronal and microglial expression patterns (Blackmore et al., 2017; Ramsden et al., 2005). Since human brain RNA-seq can only be evaluated at the time of death, there are significant challenges to resolve age-dependent changes or to definitively establish links with clinical-pathologic progression. Collection of AD mouse RNA-seq from additional timepoints may therefore accelerate discovery of improved molecular biomarkers of AD progression, and ultimately pinpoint critical windows for therapeutic interventions. Our cross-species approach additionally highlights sex-dependent changes, suggesting that female mice manifest a more rapid progression based on gene expression profiles, consistent with prior observations in both mice (Jiao et al., 2016) and human epidemiology (Altmann et al., 2014; Li and Singh, 2014; Mayeux and Stern, 2012). The apparent age by sex interaction recapitulated by our human-mouse comparisons is consistent with the companion report by Logsdon et al. (submitted), and underscores the substantial impact of sex on the brain transcriptome in AD.

Aging is the strongest known AD risk factor, and as highlighted above, is also a potent modifier of the transcriptome in AD mouse models. Using a distinct analytic design and largely independent datasets, Hargis and Blalock (2016) reported that DEGs were concordant between aging in humans and rodent models, a conclusion supported by our analysis. Strikingly, the majority of AD-associated human brain coexpression modules showing overlaps with *APP* and/or *MAPT* transgenic mouse models, were also seen in aged, wild-type mice, as well as many other disease models. This result suggests that many human brain gene expression changes associated with AD, including neuronal- and microglial-enriched modules reported in other studies (Mostafavi et al. 2018; Zhang et al. 2013), may represent common transcriptional programs activated by the aging process itself. Rather than representing a specific signature of AD pathophysiology (e.g. Aß- or Tau-mediated mechanisms), these pathways appear to be triggered and/or accelerated by heterogeneous triggers, including those manipulated in mouse models of HD, ALS, SCA1 and other neurodegenerative disorders. Interestingly, several module overlap patterns still revealed possible disease-specific signatures. For example, several modules enriched for oligodendroglial markers showed comparatively selected overlap AD models, particularly APP transgenic models, and related human brain coexpression networks have previously been implicated in AD in multiple studies (Allen et al. 2018; McKenzie et al. 2017; Mostafavi et al. 2018). Alternatively, coincident activation of both neuronal- and microglial-enriched modules was seen preferentially in models characterized by significant Tau pathologic burden. In contrast to AD and other neurodegenerative disease models, nearly all differential expression signatures from HD mice (35 out of 37) did not significantly overlap with microglial-enriched coexpression modules (Figure S4).

Our study has several notable limitations. The large size of most human coexpression modules may limit sensitivity to detect significant overlaps using the hypergeometric test, especially for functional pathways represented by smaller gene sets. Compared to the laboratory preparation of mouse mRNA, the extraction and processing of human brain tissue is likely more susceptible to postmortem artefact, although AMP-AD analyses adjusted for sample variability in postmortem interval (Logsdon et al., submitted). Moreover, the human RNA-seq data, along with the majority of included mouse studies, derive from bulk brain tissue, which includes mixed cell types. Indeed, most of the AMP-AD coexpression modules are strongly enriched for cell-type specific signatures, which may therefore reflect global changes in cell proportions, such as neuronal loss or microglial infiltration. We anticipate that the increasing availability of single cell expression profiles will definitively address this concern. In addition, since the extent and tempo of neurodegeneration, including in both human and mouse models, can vary widely across different brain regions, RNA-seq profiles from whole brain might obscure more localized transcriptome overlaps. Lastly, while we selected nearly 100 independent mouse RNAseq studies for inclusion in our analyses, prioritizing those most relevant to AD and neurodegeneration, we omitted many others with the potential to provide additional insights. In the future, our approach can thus be generalized to examine overlaps with an even broader sample of available mouse data.

The strengths of our study include consideration of a still large and diverse number of mouse studies and the use of a single RNAseq-reprocessing pipeline to facilitate cross-model comparisons. We also leveraged AMP-AD coexpression modules based on analyses of more 2,000 human brains and representing consensus networks derived from 5 independent algorithms. Importantly, unlike comparisons based on pathology or behavioral phenotypes, brain expression profiles likely represent more proximal endophenotypes, potentially affording greater sensitivity and reliability for detection of cross-species overlaps. In fact, we highlight several overlaps with transcriptomic endophenotypes from completely unexpected mouse experimental manipulations which manifest brain expression changes that mimic human AD, and in some cases even overlap modules better than currently available AD mouse models. Such “AD transcriptologs”—mouse models based on transcriptome homology—may pinpoint non-obvious experimental models for future investigation of AD pathophysiology.

## METHODS

### Human coexpression modules

The derivation of 30 AD-associated human coexpression modules is described in the companion report from AMP-AD (Logsdon et al., submitted) (doi: 10.7303/syn11932957.1). Briefly, the consensus networks are based on RNA-Seq data from 2114 total postmortem brain samples from 3 independent cohorts, including the Religious Order Study and the Memory and Aging Project (syn3388564) (Bennett et al., 2018; Mostafavi et al., 2018), the Mount Sinai Brain Bank (syn3157743) (Wang et al., 2018), and the Mayo clinic (syn5049298, syn3163039) (Allen et al., 2016; Allen et al., 2018). Postmortem tissue was collected from seven distinct brain regions: dorsolateral prefrontal cortex (DLPFC); temporal cortex (TCX) and cerebellum (CBE), and inferior frontal gyrus (IFG), superior temporal gyrus (STG), frontal pole (FP), and parahippocampal gyrus (PHG). Five distinct algorithms were used to independently generate coexpression networks from each brain region, and graph clustering was subsequently applied to all modules enriched for AD-differentially expressed genes, leading to 30 aggregate modules (Table S1).

### Mouse dataset selection

Figure 1 depicts the overall analysis pipeline for collecting and processing mouse studies, and examining overlaps with human coexpression modules. A total of 96 studies, encompassing data from 2279 mouse tissue samples, were analyzed (Table S2). These data were collected from three sources: the Gene Expression Omnibus (GEO) database (83 studies), the AMP-AD Knowledge portal (syn5550383) (6 studies), and through personal communication (7 studies). We searched the GEO database on September 12, 2017 using the keywords “brain”, “mouse”, and “expression profiling by high throughput sequencing”, identifying 881 studies for initial consideration. All studies were next indexed using high frequency terms, and secondary filtering was based on manually curated keywords relevant to AD pathophysiology (Karch et al., 2014) (Table S2). Lastly, the filtered list of 349 studies was reviewed by the study team. We excluded studies, or in some cases specific samples, involving (1) tissue source other than nervous system, (2) organisms other than mouse, or (3) non-coding RNA. From the remainder, 79 GEO studies were selected for inclusion based on relevance to AD, neurodegeneration, or related mechanisms. All included studies had publicly available RNA-seq data derived from either mouse brain or brain-derived cell lines. Following these initial searches, we discovered only a single eligible RNA-seq study for analysis of aging-associated expression changes (GSE61915). Given the importance of aging in AD, we identified and included an additional 5 expression array profiling studies related to brain aging. Table S2 details all studies included in this analysis, including data source, citations, and relevant keywords.

### RNA-seq/microarray re-processing and differential expression analysis

A unified RNA-seq analysis pipeline (Figure 1) was used for reprocessing of all datasets, with the exception of 2 HD studies where count files were downloaded directly from GEO. Data processing leveraged the cloud formation cluster at Amazon Web Services. We first created one EC2 master instance (m3.xlarge) which was used to launch hundreds of EC2 computing nodes (c3.8xlarge). Next, each computing node was assigned to process one sample using our customized RNA-seq pipeline, as implemented using Snakemake v.4.8.0 (Köster and Rahmann, 2012). For samples in AMP-AD studies, the pipeline begins with downloading BAM files from the AMP-AD Knowledge Portal using the Synapse python client. The BAM files are then converted to fastq files using Picard SamToFastq command v.2.18.2 (http://broadinstitute.github.io/picard). For GEO studies, SRA files were downloaded from the database using the GEOquery R package v.2.42.0 (Davis and Meltzer, 2007), and fastq files were generated using the fastq-dump command from the NCBI SRA toolkit v2.8.2.1 (Andrews, 2010). Alignment to the mouse reference genome GRCm38 (mm10) was implemented using STAR v.2.5.1b (Dobin et al., 2013), and BAM file reads were subsequently sorted by coordinate using samtools v0.1.5 (Li et al., 2009). Genes were quantified using either HTSeq v0.6.0 (Anders et al., 2015) or using the ‘quantMode’ option from the STAR aligner which utilizes HTSeq algorithm and produces similar results. Results were uploaded to the Synapse portal using the python client. Differential gene expression analysis was conducted using DESeq2 v1.18.1 (Love et al., 2014). For the limited number of microarray studies, pre-processed intensities available from the series matrix files were downloaded from GEO and normalized using quantile normalization, followed by differential expression analysis using the limma package v3.4.2 (Ritchie et al., 2015). For each study (Table S3), experimental and control pairs were manually curated. We required a minimum of n = 2 samples for each group (experimental and control); the average for all samples included in each comparison was n = 8.4 (range = 4-28 total samples). Overall, 376 mouse experimental comparisons were curated for computation of differentially expressed gene sets (DEGs), applying a false-discovery rate (FDR) threshold of 1% and minimum fold-change of 1.2. The t-distributed Stochastic Neighbor Embedding (t-SNE) algorithm (Maaten and Hinton, 2008) was applied to DEGs from all studies (logarithm-transformed fold-change), using the Rtsne function in R. We excluded all DEG sets from consideration consisting of fewer than 10 conserved mouse genes, resulting in 251 sets of DEGs for consideration in our subsequent analyses. All analysis was done using R v3.4.2, and Python v.2.7.12. Table S3 details all DEG sets meeting these criteria, including data sources.

To facilitate ease of use and repurposing of the data, each mouse DEG set was assigned a unique identifier (M###), and we also developed a descriptive nomenclature for annotation. Each gene set received a label taking the form: *category_experimental condition_sex_age_brain region_cell type_ transgene*. In this standardized annotation, “category” denotes the relevant neurologic disease (e.g. AD, HD, SCA, ALS) or “other” for gene manipulations not directly linked to human disease, along with the specific “experimental condition” describing the mouse genotype or treatment condition. We also note “sex” (M or F), “age” (months), and where applicable, “brain region” (e.g. hippocampus), “cell type” (e.g. neuron, microglia). In the case of AD mouse models, we also annotate “transgene”, to differentiate “APP”, “Tau” (MAPT), or “other” models. If unknown or not applicable, the relevant field(s) are replaced with “na”. These annotations and conventions are used throughout our supplementary tables and files.

### Analysis of mouse-human overlaps

Mouse orthologs for all human genes were extracted using the HCOP tool available from the HUGO Gene Nomenclature Committee (https://www.genenames.org/cgi-bin/hcop) (syn17010253). Using the hypergeometric test, we determined the significance of overlap between each of 251 mouse DEG sets (above) and the 30 human gene coexpression modules (mouse orthologs) using the phyper function in R (Table S4). The Benjamini-Hochberg method was applied to adjust for multiple comparisons, using the p.adjust function. All p-values reported in the text were adjusted in this manner. Overlap significance was visualized using heatmaps, implemented with pheatmap function in R, using Manhattan distance (for both rows and columns) and Ward clustering. In order to determine whether mouse-human overlapping genes also shared expression changes in the same direction, we computed the concordance score for each overlap. The direction of change for human genes was extracted from the data in Logsdon et al. (submitted) (syn14237651). Specifically, the concordance score is the percentage of genes in the concordant direction weighted by the significance based on the hypergeometric test, which is computed as follows:

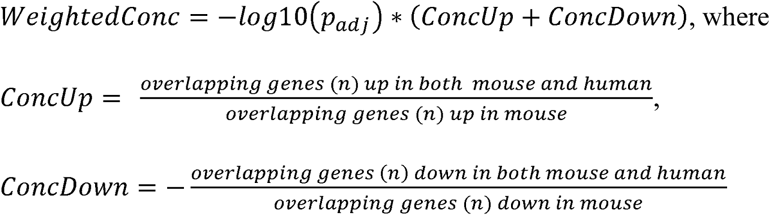

The overlap was considered concordant when the weighted concordance is greater (or less) than half standard deviation from the median, and p_adj_ ≤ 0.01 for the up- or down-differential expression.

## ACKNOWLEDGEMENTS

This study was supported in part by grants from the NIH (R01AG053960, R01AG050631, R01AG057339, U01AG046161, U01 AG046139, U01AG046170, U01AG046152, P30AG10161, R01AG15819, R01AG17917, R01AG057473, R01AG057914, P50 NS38377, F31AG050357, P50 AG016574, R01 AG032990, R01 AG018023, U01 AG006576, U01 AG006786, R01 AG025711, R01 AG017216, R01 AG003949, R01 NS080820, U24 NS072026, P30 AG19610). J.M.S. was additionally supported by grants from Huffington Foundation, Jan and Dan Duncan Neurological Research Institute at Texas Children’s Hospital, and a Career Award for Medical Scientists from the Burroughs Wellcome Fund. Z.L. received additional support from the National Science Foundation - Division of Mathematical Sciences DMS-1263932, Cancer Prevention Research Institute of Texas RP170387, Houston Endowment and the Neurodegeneration Consortium and Belfer Family Foundation. T.M.D. is supported by the JPB Foundation and is the Leonard and Madlyn Abramson Professor in Neurodegenerative Diseases. The authors acknowledge the joint participation by the Adrienne Helis Malvin Medical Research Foundation through its direct engagement in the continuous active conduct of medical research in conjunction with The Johns Hopkins Hospital and the Johns Hopkins University School of Medicine and the Foundation’s Parkinson’s Disease Program M-2014. Mayo Clinic RNAseq Study were additionally supported by the Mayo Clinic Alzheimer’s Disease Genetic Studies, led by Dr. Nilufer Ertekin-Taner and Dr. Steven G. Younkin, Mayo Clinic, Jacksonville, FL using samples from the Mayo Clinic Study of Aging, the Mayo Clinic Alzheimer’s Disease Research Center, and the Mayo Clinic Brain Bank; CurePSP Foundation; Mayo Foundation; the Sun Health Research Institute Brain and Body Donation Program of Sun City, Arizona; the Brain and Body Donation Program; Arizona Alzheimer’s Disease Core Center; the Arizona Department of Health Services (contract 211002, Arizona Alzheimer’s Research Center), the Arizona Biomedical Research Commission (contracts 4001, 0011, 05-901 and 1001 to the Arizona Parkinson’s Disease Consortium) and the Michael J. Fox Foundation for Parkinson’s Research.

We are grateful to Dr. Bruce Yankner for contributing additional mouse RNA-sequencing data. This study is part of the National Institute on Aging Accelerating Medicines Partnership-AD, MODEL-AD, and Resilience-AD programs. Data in the AMP-AD Knowledge Portal that contributed to or was generated from this study can be accessed at doi:10.7303/syn2580853.

## AUTHOR CONTRIBUTIONS

Conceptualization, Y.-W.W., Z.L., J.M.S.; Methodology, Y.-W.W., R.A.-L., Z.L.; Data Curation, R.A.-L., T.V.L., K.A.; Formal Analysis, Y.-W.W., R.A.-L., C.G.M., B.L.; Writing—Original Draft, Y.-W.W., C.G.M., R.A.-L., T.V.L., J.M.S.; Writing—Review & Editing, C.G.M., R.A.-L., J.M.S., Z.L., T.V.L., K.A., S.N., C.K., H.Z., V.D., A.I.L., B.L., L.M.; Funding Acquisition, J.M.S., Z.L., L.M., A.L., T.A.G.; Resources, S.N., C.K., V.P., G.H., H.M.S., H.Z., J.W.K., V.D., T.D., P.-C.P., L.-H.T., J.-V.H.-M., M.W., M.E.E., H.M., X.Z., P.C., Y.L., T.A.G., B.L., L.M.; Supervision, J.M.S., Z.L.

## DATA AND SOFTWARE AVAILABILITY

Data from this study, including all re-processed RNA-seq from mouse models, subsequent differential expression analysis, and the overlap analyses with human coexpression modules is available from the AMP-AD Knowledge Portal at doi:10.7303/syn16779040.

## SUPPLEMENTAL INFORMATION

Supplemental Information includes 6 figures and 3 tables.

