## Supplemental Information for "Functional dissection of Alzheimer’s disease brain gene expression signatures in humans and mouse models"

### SUPPLEMENTAL FIGURES

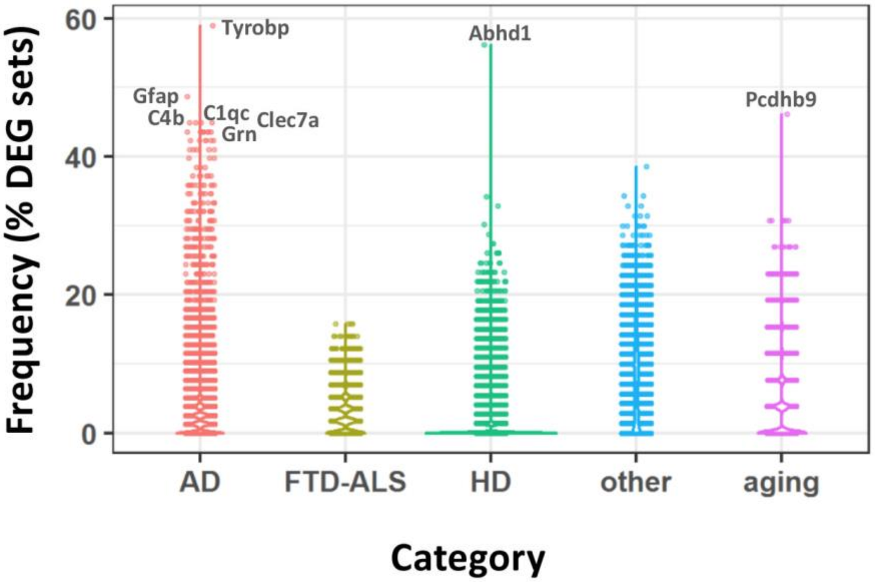


**Figure S1: Mouse expression signatures are highly heterogeneous (Related to Figure 1).**
Recurrence rates for individual genes are shown among mouse DEG sets within each disease category. Most DEGs appear in only a minority of expression signatures, for a given disease category. Several highly recurrent genes among AD DEG sets are noted, several of which have roles in inflammation and innate immunity (see also Figure S3).

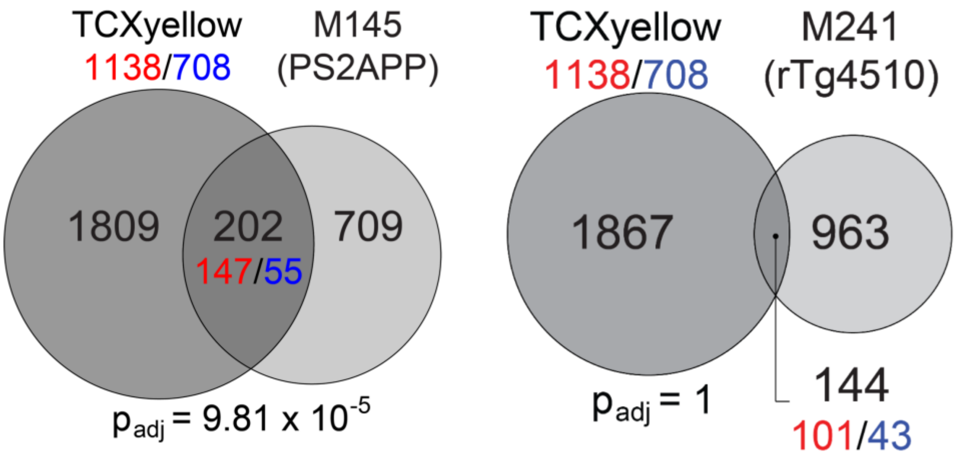


**Figure S2: Module overlap specificity for APP expression signatures (Related to Figure 3).**
Representative overlaps for the human TCXyellow coexpression module, showing relative specificity for shared gene expression changes with APP (LEFT: M145, PS2APP) but not Tau (RIGHT: M241, rTg4510) signatures. Gene counts are noted in black, including for overlapping and non-overlapping regions. To assess concordance between human brains and mouse models, gene counts are shown, noting increased or decreased expression (red or blue, respectively), including for the whole human coexpression module and the overlapping gene set from mouse models. TCXyellow significantly overlaps with 2 out of 26 APP expression signatures, but not with any MAPT or other AD models.

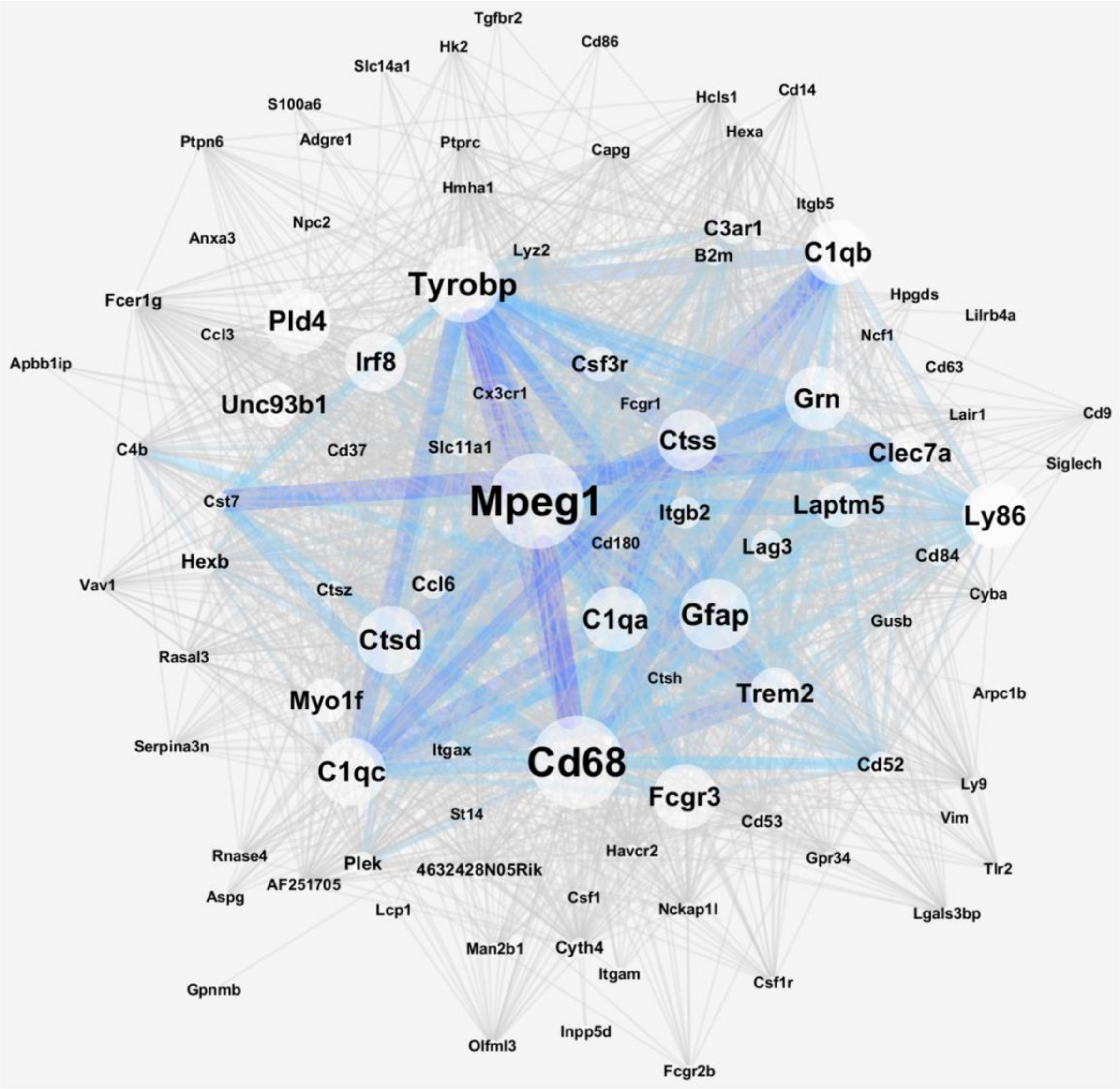


**<u>Figure S3: Commonly recurring mouse DEGs comprise an inflammation expression network (Related to Figure 1).</u>**
Network depicting the most highly recurrent genes across 251 mouse model DEG sets and their co-occurrence relationships. Edges indicate co-occurrence of genes within a given model DEG set, with edge width indicating the number of times these genes recur together. The 95 genes depicted occur in at least 10% of the 376 DEG sets included in this study, and most significantly enriched for genes implicated in inflammation and immune system processes (GO:0002376, FDR=4.76x$10^{-10}$). Gene-pair co-occurrence counts were computed in R, and the network was constructed using the igraph package and visualized with Cytoscape. DAVID Bioinformatics Resources 6.8 functional tools was used to determine functional enrichment based on Gene Ontology (GO) terms.

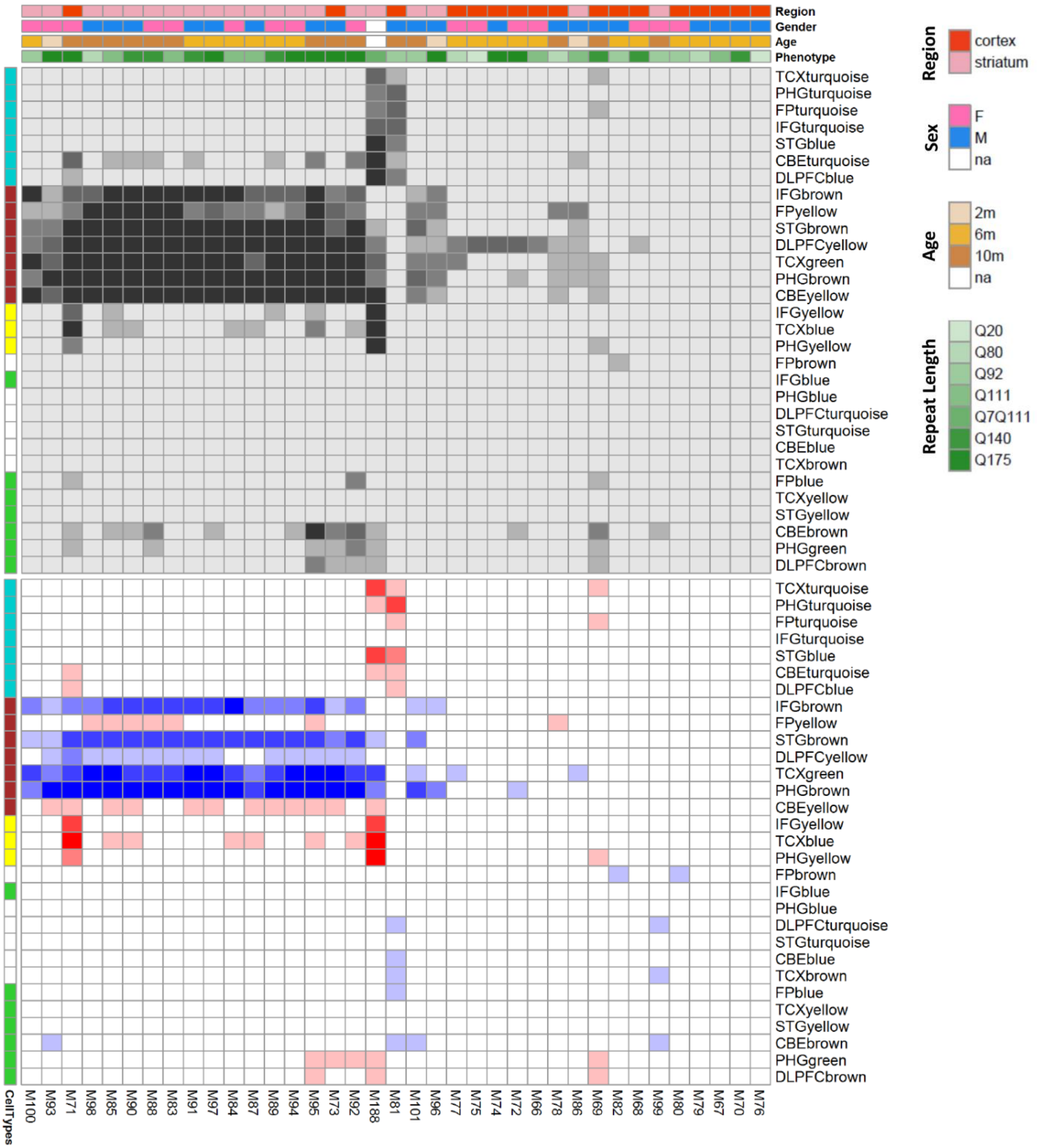


**Figure S4: Additional overlap with Huntington's disease models (Related to Figure 4).**

Heat maps show overlap (top) and concordance (bottom) among 30 human coexpression modules and sets of differentially expressed genes (DEGs) derived from Huntington's disease (HD) mouse models. Mouse-human overlap significance, calculated using the hypergeometric test, is represented in grayscale [$-\log_{10}(p_{adj})$]. Annotation of direction (red/blue) and concordance (intensity) for mouse/human gene expression changes, as well as human module cell type enrichment following conventions in Figure 4.

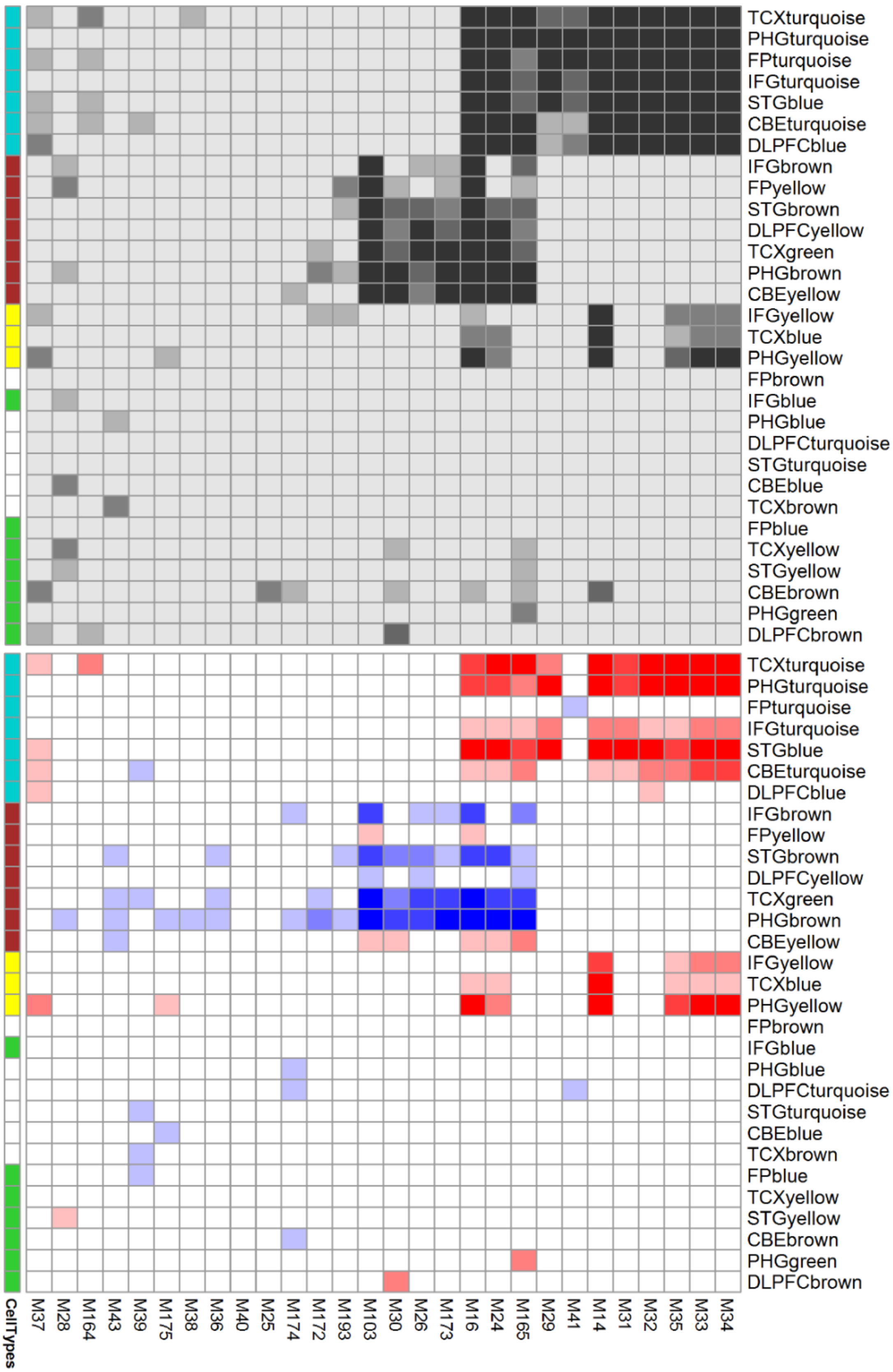


**Figure S5: Additional overlap with FTD-ALS models (Related to Figure 4).**

Heat maps show overlap (top) and concordance (bottom) among 30 human coexpression modules and sets of differentially expressed genes (DEGs) derived from frontotemporal dementia-amyotrophic lateral sclerosis (FTD-ALS) mouse models. Mouse-human overlap significance, calculated using the hypergeometric test, is represented in grayscale [$-\log_{10}(p_{adj})$]. Annotation of direction (red/blue) and concordance (intensity) for mouse/human gene expression changes, as well as human module cell type enrichment following conventions in Figure 4.

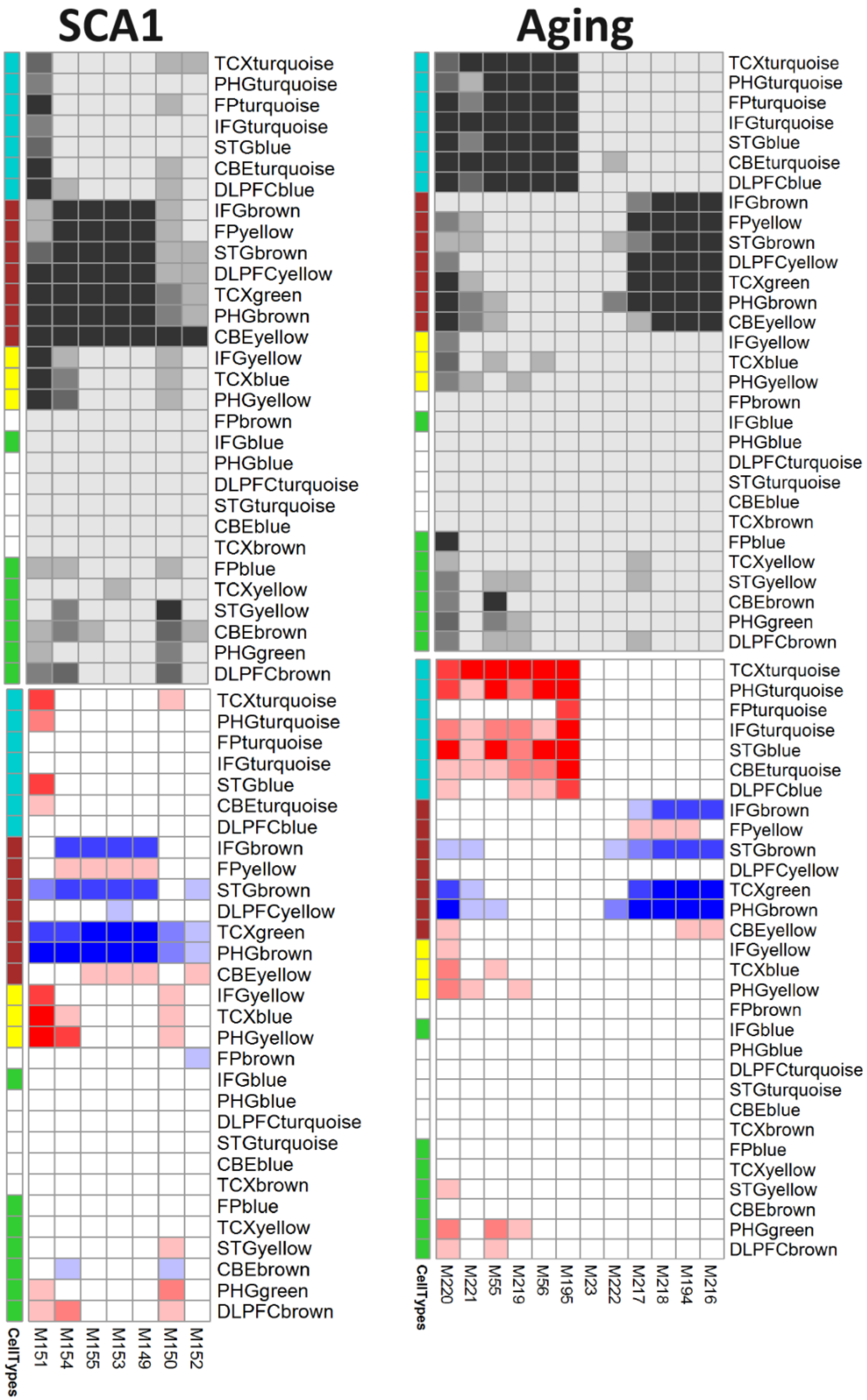


**Figure S6: Additional overlap with SCA1 and aging models (Related to Figure 4).**

Heat maps show overlap (top) and concordance (bottom) among 30 human coexpression modules and sets of differentially expressed genes (DEGs) derived from spinocerebellar ataxia, type 1 (SCA1, Left) and aging mouse models (Right). Mouse-human overlap significance, calculated using the hypergeometric test, is represented in grayscale [$-\log_{10}(p_{adj})$]. Annotation of direction (red/blue) and concordance (intensity) for mouse/human gene expression changes, as well as human module cell type enrichment following conventions in Fig. 4.
